# Diverse pathogens activate the host RIDD pathway to subvert BLOS1-directed immune defense

**DOI:** 10.1101/2021.11.04.467275

**Authors:** Kelsey Wells, Kai He, Aseem Pandey, Ana Cabello, Dong-Mei Zhang, Jing Yang, Gabriel Gomez, Yue Liu, Hao-Wu Chang, Xue-Qing Li, Hao Zhang, Luciana Fachini da Costa, Richard P. Metz, Charles D. Johnson, Cameron Martin, Jill Skrobarczyk, Luc R. Berghman, Kristin Patrick, Julian Leibowitz, Allison Rice-Ficht, Sing-Hoi Sze, Xiaoning Qian, Qing-Ming Qin, Thomas A. Ficht, Paul de Figueiredo

## Abstract

The phagocytosis and destruction of pathogens in lysosomes constitute central elements of innate immune defense. Here, we show that *Brucella*, the causative agent of brucellosis, the most prevalent bacterial zoonosis globally, subverts this immune defense pathway by activating regulated IRE1α-dependent decay (RIDD) of mRNAs encoding BLOS1, a protein that promotes endosome-lysosome fusion. RIDD-deficient cells and mice harboring a RIDD-incompetent variant of IRE1α were resistant to infection. Non-functional *Blos1* struggled to assemble the BLOC-1-related complex (BORC), resulting in differential recruitment of BORC-related lysosome trafficking components, perinuclear trafficking of *Brucella*-containing vacuoles (BCVs), and enhanced susceptibility to infection. The RIDD-resistant *Blos1* variant maintains the integrity of BORC and a higher-level association of BORC-related components that promote centrifugal lysosome trafficking, resulting in enhanced BCV peripheral trafficking and lysosomal-destruction, and resistance to infection. These findings demonstrate that host RIDD activity on BLOS1 regulates *Brucella* intracellular parasitism by disrupting BORC-directed lysosomal trafficking. Notably, coronavirus MHV also subverted the RIDD-BLOS1 axis to promote intracellular replication. Our work therefore establishes BLOS1 as a novel immune defense factor whose activity is hijacked by diverse pathogens.

## INTRODUCTION

*Brucella* is an intracellular vacuolar pathogen that invades many cell and tissue types, including non-professional and professional phagocytes (de Figueiredo *et al*, 2015). Brucellosis has eluded systematic attempts at eradication for more than a century (Godfroid *et al*, 2002), and even in most developed countries, no approved human vaccine is available (Ficht & Adams, 2009). The intracellular lifestyle limits exposure to host innate and adaptive immune responses and sequesters the organism from the effects of some antibiotics. *Brucella* evades intracellular destruction by limiting interactions of the *Brucella*–containing vacuole (BCV) with the lysosomal compartment (Criscitiello *et al*, 2013; Pizarro-Cerda *et al*, 1998). BCVs harboring internalized *Brucella* traffic from endocytic compartments (eBCVs) to a replicative niche within vacuoles (rBCVs) that are decorated with markers of the endoplasmic reticulum (ER) (Pizarro-Cerda *et al*., 1998; Starr *et al*, 2012). BCVs also accumulate autophagic membranes (aBCVs), which constitute a distinctive aspect of the intracellular lifestyle of the pathogen (Pandey *et al*, 2018; Starr *et al*., 2012). The VirB type IV secretion system (T4SS) is a significant virulence factor that regulates *Brucella* intracellular trafficking (Marchesini *et al*, 2011; Paredes-Cervantes *et al*, 2011; Sa *et al*, 2012; Smith *et al*, 2012). *Brucella* effectors secreted by the T4SS promote bacterial intracellular trafficking and growth via modulation of host functions (de Barsy *et al*, 2011; De Jong *et al*, 2008; Dohmer Pisani *et al*, 2014; Miller *et al*, 2017; Myeni *et al*, 2013) and organisms that lack this system fail to establish productive infections.

The Unfolded Protein Response (UPR) is an evolutionarily conserved signaling pathway that allows the ER to recover from the accumulation of misfolded proteins (Gardner *et al*, 2013; Walter & Ron, 2011) during ER stress. The UPR signals through the stress sensors IRE1α, ATF6, and PERK located in the ER membrane. When the luminal domains of these proteins sense unfolded proteins, they transduce signals to their cytoplasmic domains, which initiate signaling that ultimately results in UPR (Lee *et al*, 2008). IRE1α plays a central role in triggering UPR through an endonuclease/RNase activity in its cytoplasmic tail that catalyzes the splicing of *Xbp1* mRNA, which is then translated to generate the XBP1 transcription factor (Lee *et al*., 2008; Ron & Walter, 2007). IRE1α RNase activity can also cleave a wide variety of cellular mRNAs that leads to their degradation in a process termed regulated IRE1-dependent mRNA decay (RIDD) (Hollien & Weissman, 2006). The RIDD pathway displays selectivity. For example, the pathway cleaves a specific subset of mRNAs encoding polypeptides destined for co-translational translocation into the ER lumen. The degradation of these mRNAs supports ER homeostasis by reducing the flux of non-essential polypeptides into the ER (Hollien & Weissman, 2006). The molecular targets of RIDD activity, and the physiological roles that this process plays in cells, remain areas of investigation.

*Brucella* infection induces host cell ER stress and activates host UPR (de Jong *et al*, 2013; Pandey *et al*., 2018; Smith *et al*, 2013; Taguchi *et al*, 2015; Wang *et al*, 2016). The UPR sensor IRE1α, but neither PERK nor ATF6, is required for the intracellular replication of the pathogen (Qin *et al*, 2008; Taguchi *et al*., 2015), indicating that the IRE1α signaling pathway confers susceptibility to host cell parasitism. An IRE1α-ULK1 signaling axis also contributes to conferring susceptibility to *Brucella* intracellular replication; IRE1α-directed activation of components of the host autophagy program promotes proper bacterial intracellular trafficking and replication (Pandey *et al*., 2018). Despite the above-mentioned advances, our understanding of how the IRE1α-RIDD axis and downstream processes regulate the intracellular lifestyle of *Brucella* remains largely unknown.

BLOS1 [biogenesis of lysosome-related organelles complex-1 (BLOC-1) subunit 1, also known as BLOC1S1/GCN5L1], a subunit of both the BLOC-1 and the BLOC-1-related complex (BORC), plays diverse roles in cells, including mitochondrial protein acetylation, modulation of metabolic pathways, and endosome-lysosome trafficking and fusion (Bae *et al*, 2019; Guardia *et al*, 2016; Pu *et al*, 2017; Pu *et al*, 2015). In mammalian cells, BLOS1 has also been shown to be a principal target of RIDD activity (Bae *et al*., 2019; Bright *et al*, 2015; Hollien *et al*, 2009) and is required for host cell cytotoxicity induced by Ebola virus (Carette *et al*, 2011). To date, the roles and mechanisms by which BLOS1 controls infection by intracellular pathogens remain largely unknown. Here, we demonstrate that IRE1α-directed *Blos1* mRNA degradation confers susceptibility to *Brucella* infection. *Brucella***-** induced RIDD activity suppresses *Blos1* expression, disassembles BORC components, and limits BLOS1-regulated interactions between BCVs and lysosomes. In addition, we show that murine hepatitis virus (MHV), a betacoronavirus, also subverts BLOS1 activity to promote its intracellular replication. Collectively, these activities promote the productive subcellular trafficking and intracellular replication of diverse pathogens. Our findings, therefore, identify BLOS1 as a novel immune defense factor that defends against bacterial and viral infection and show that *Brucella* and MHV subvert this innate immune defense system to promote disease.

## Results

### IRE1*α* conditional knockout (CKO, LysM-IRE1*α*^-/-^) mice are resistant to *Brucella* infection

*Brucella* induces host cell UPR during infection (Qin *et al*., 2008; Taguchi *et al*., 2015) and activates an IRE1α-to-autophagy signaling axis in host cells to promote its intracellular lifestyle (Pandey *et al*., 2018). To extend these findings to an *in vivo* model of brucellosis, we tested the hypothesis that UPR and IRE1α confer susceptibility to *Brucella* infection in mice harboring a conditional mutation in *Ern1*, the gene encoding IRE1α. Because mice homozygous for null mutations in *Ern1* display embryonic lethality during organogenesis, we generated a control LysM-IRE1α ^wt/wt^ [wt (wild-type)-IRE1α] and a IRE1α CKO mouse line, LysM-IRE1α ^mut/mut^ (m-IRE1α) (**Figure 1—figure supplement 1A)**. In this line, exons 20-21 of the gene encoding IRE1α were deleted in monocytes and macrophages, generating animals in which the endonuclease domain (and hence RIDD activity) was specifically disrupted. However, the kinase domain remained intact (Hur *et al*, 2012; Iwawaki *et al*, 2009). Macrophages are critical cellular targets for *Brucella* colonization (de Figueiredo *et al*., 2015). Hence, the tissue and molecular specificity of this lesion rendered the m-IRE1α mouse an ideal system for investigating how bacterial activation of host RIDD activity controls intracellular parasitism by the virulent *B. melitensis* strain 16M (Bm16M).

After confirming the RNase activity deficiency of the truncated IRE1α in bone marrow-derived macrophages (BMDMs) from m-IRE1α animals (**Figure 1—figure supplement 1B**), we verified that m-IRE1α mice had normal organ morphologies, fertility, growth, and development. In addition, we showed that these animals had similar B cell (B220^+^), T cell (CD4^+^ or CD8^+^), and CD11b^+^ profiles (**Figure 1—figure supplement 1C**). We also showed that the IL-1β, IL-6 and TNF-α responses of BMDMs to LPS stimulation were reduced in mutant mice (**Figure 1A-C**), consistent with previous findings that these LPS-mediated responses are controlled, in part, by IRE1α activity (Martinon *et al*, 2010).

**Figure 1.**
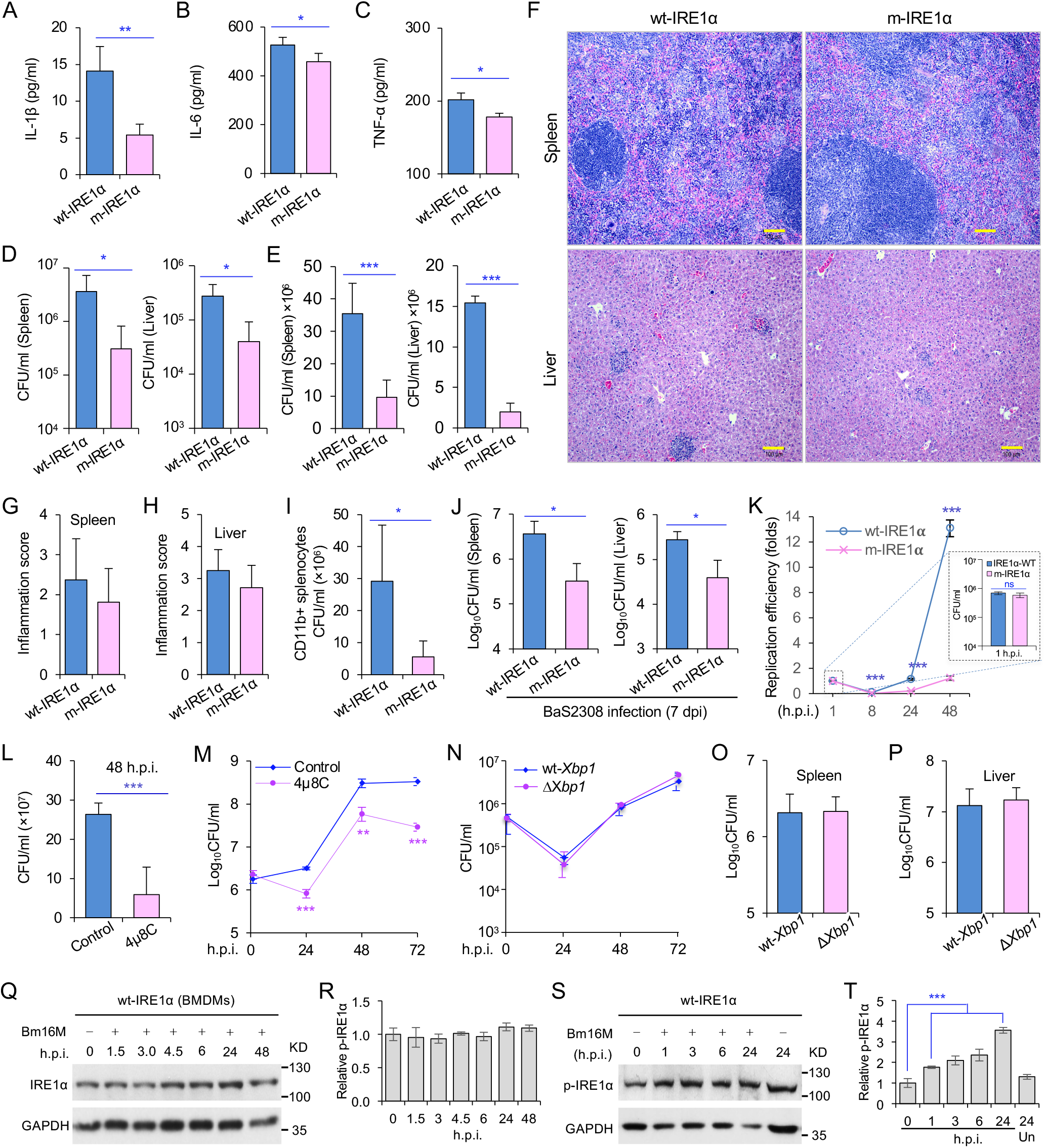
Host IRE1*α* is required for *Brucella* infection *in vivo*. (A-C) Innate cytokine production of IL-1 β (A), IL-6 (B), and TNF-α (C) in brow marrow derived macrophages (BMDMs) from the wild-type (WT, wt-IRE1α) control and *IRE1α* conditional knockout (CKO, m-IRE1α) mice. (D, E) Colony-forming unit (CFU) assay for *B. melitensis* 16M (Bm16M) intracellular survival in spleens and livers of wt- and m-IRE1α mice at 7 (D) or 14 (E) days post infection (dpi).(F) Histopathology of representative hematoxylin and eosin (H&E) stained sections of spleen and liver from Bm16M infected wt- and m-IRE1α mice at 14 dpi. Bars: 100 μm.(G, H) Quantification of inflammation of spleens (G) or livers (H) at 14 dpi.(I) CFU assays of CD11b^+^ cells from Bm16M infected wt- or m-IRE1α mice.(J) CFU assay for *B. abortus* S2308 (BaS2308) intracellular survival in spleens and livers in wt-IRE1α control or m-IRE1α mice at 7 dpi.(K) Bm16M invasion (right inset) and intracellular replication in BMDMs from m-IRE1α and control mice. h.p.i.: hours post infection.(L, M) CFU assays of Bm16M infection of WT BMDMs (L) or RAW264.7 macrophages (M). Host cells were pretreated with 4μ8C (50 μM) 1 hr before and during infection; CFUs of the infected cells were determined at the indicated h.p.i..(N) CFU assays for Bm16M infection of BMDMs from WT and *Xbp1* knockout (Δ*Xbp1*) mice at the indicated h.p.i..(O, P) CFU assay for Bm16M intracellular survival in spleen (O) or liver (P) in WT or Δ*Xbp1* mice at 14 dpi.(Q, R) Immunoblotting assay for IRE1α expression (Q) and quantification of the expression levels (R) in BMDMs during a time course (48 hr) of Bm16M infection.(S, T) Bm16M infection induces phosphorylation of host IRE1α (S) and quantification of the phosphorylated levels of IRE1α during a time course (24 hr) of infection (T).Images/blots are representative of three independent experiments. Statistical data represent the mean ± SEM (standard error of mean) from three independent experiments. *, **, *** indicates significance at *p* < 0.05, 0.01 and 0.001, respectively. **Figure 1—figure supplement 1**. Characterization of IRE1α conditional knockout (CKO) and control mice. **Figure 1—figure supplement 2**. IRE1α is required for *B. melitensis* intracellular replication.

To determine whether IRE1α activity in macrophages contributed to pathogen burden, dissemination, and disease progression, we infected wt-IRE1α control and m-IRE1α mice with Bm16M via the intraperitoneal route, humanely sacrificed the mice at various times post-infection, and then determined the bacterial burden in assorted tissues by quantifying the number of recovered colony forming units (CFUs). We found that tissue-specific mutation of IRE1α resulted in enhanced resistance to bacterial infection with significant reductions in bacterial load in the spleen and liver compared to wt-IRE1α controls at 7 or 14 days post-infection (dpi), respectively (**Figure 1D-E**). However, both infected wt- and m-IRE1α mice displayed similar spleen weights (**Figure 1—figure supplement 1D-E**) and spleen or liver inflammation (**Figure 1F-H**), revealing that the lower numbers of CFU recovered from m-IRE1α animals were not accompanied by corresponding decreases in inflammation. To test the hypothesis that the differential bacterial burden in macrophage cells, the predominant cell type in which the pathogen resides and replicates *in vivo*, accounted for this reduction, we compared the bacterial load in CD11b^+^ cells from control and m-IRE1α mice that had been infected with Bm16M for seven days. We found that indeed CD11b^+^ cells from the spleens of m-IRE1α mice displayed striking reductions in bacterial load (**Figure 1I**), thereby suggesting that the resistance of these cells to intracellular parasitism contributed to the resistance phenotype observed at the organismal level. Divergent *Brucella* species display distinct host preferences; however, their interactions with host cells share common features (de Figueiredo *et al*., 2015). To test the hypothesis that m-IRE1α mice also displayed resistance to infection by other *Brucella* species, we infected these mice with *B. abortus* strain S2308 (BaS2308), a strain that displays tropism for cattle. We then assessed tissue burden in spleen and liver at 7 dpi. We found that m-IRE1α mice also exhibited resistance to BaS2308 infection (**Figure 1J**), thereby indicating that the resistance phenotype of the mutant mice was not pathogen species-specific.

### IRE1*α* RNase activity confers susceptibility to *Brucella* infection

*Xbp1* splicing was dramatically diminished in BMDMs from m-IRE1α mice (**Figure 1—figure supplement 1B)**, indicating that BMDMs from the m-IRE1α mice carried the expected functional defects in IRE1α RNase activity. We thus tested the hypothesis that IRE1α RNase activity confers susceptibility to intracellular parasitism by Bm16M. First, we performed CFU assays of Bm16M infection of BMDMs from m-IRE1α and control mice and found that the replication efficiency of Bm16M in m-IRE1α BMDMs at 8, 24, and 48 h.p.i. was significantly lower than controls (**Figure 1K**). Similar results were observed in control and IRE1α^-/-^ mouse embryonic fibroblasts (MEFs) (**Qin et al**., **2008; Figure 1—figure supplement 2A-B**). Second, we observed fewer Bm16M were recovered from BMDMs or RAW264.7 macrophages treated with 4μ8C, a compound that specifically antagonizes IRE1α RNase activity while leaving its kinase activity intact (Cross *et al*, 2012), than mock-treated controls (**Figure 1L-M**). These findings supported the hypothesis that IRE1α RNase activity confers susceptibility to infection by regulating the intracellular trafficking and/or survival of the pathogen.

### IRE1*α* regulation of Bm16M intracellular replication is XBP1-independent

We performed additional experiments to interrogate the role of IRE1α RNase activity in controlling *Brucella* infection. Two possibilities were explored: (1) IRE1α catalyzed splicing of *Xbp1* transcripts, and downstream expression of XBP1-responsive genes, conferred susceptibility to Bm16M infection; or (2) IRE1α catalyzed RIDD activity controlled this process. To determine whether *Xbp1* splicing was the reason, we examined the survival and intracellular replication of the pathogen in BMDMs harvested from LyzM-*Xbp1*^-/-^ mice in which *Xbp1* was conditionally ablated from monocytes, macrophages, and granulocytes (henceforth, Δ*Xbp1* mice) and found that Bm16M replicated similarly in Δ*Xbp1* BMDMs and wt-*Xbp1* littermate controls (**Figure 1N**). Moreover, Bm16M displayed similar levels of liver and spleen colonization in Δ*Xbp1* and wt-*Xbp1* mice (**Figure 1O-P**). These data suggested an XBP1-independent role for IRE1α RNase activity in conferring susceptibility to Bm16M infection, thereby implicating RIDD as the sought-after activity (see below).

### IRE1*α* activity regulates *Brucella* intracellular trafficking and replication

To dissect the mechanism by which IRE1α activity confers susceptibility to intracellular parasitism, we examined the intracellular trafficking and replication of Bm16M in BMDMs derived from m-IRE1α mice. We observed that the expression level of IRE1α was relatively unchanged during a time course (48 hr) of infection in both wt-IRE1α BMDMs (**Figure 1Q-R**) and MEFs (**Figure 1—figure supplement 2C**). IRE1α phosphorylation was enhanced over the same time course in wt-IRE1α BMDMs during Bm16M infection (**Figure 1S-T**). To assess the influence of host IRE1α activity on the intracellular replication of the pathogen, we used gentamicin protection analysis (Qin *et al*., 2008) to examine bacterial replication in BMDMs from control and m-IRE1α mice, as well as in control and IRE1α^-/-^ MEFs. As expected, Bm16M replicated efficiently in wt-IRE1α BMDMs and control MEFs; however, the replication efficiency was significantly diminished in m-IRE1α BMDMs and IRE1α^-/-^ MEFs (**Figure 1K; Figure 1—figure supplement 2B**). To test whether IRE1α regulates Bm16M intracellular trafficking, we used confocal immunofluorescence microscopy (CIM) to analyze the localization of the pathogen in IRE1α^+/+^ and IRE1α^-/-^ MEFs, or m-IRE1α and control BMDMs. In IRE1α harboring controls, the pathogen transiently trafficked through early and late endosomes (EEA1^+^ and M6PR^+^ compartments, respectively) (**Figure 2A-D; Figure 2—figure supplement 1A-D**) before primarily accumulating (at 24 and 48 h.p.i.) in a replicative niche decorated with the ER marker calreticulin (**Figure 2E-F; Figure 2—figure supplement 1E and G**); however, in m-IRE1α BMDMs or IRE1α^-/-^ MEF cells, Bm16M displayed reduced trafficking to calreticulin^+^ compartments (**Figure 2E-F; Figure 2—figure supplement 1E and G**). Instead, the pathogen trafficked with greater efficiency to M6PR^+^ late endosomes (at 12 h.p.i) (**Figure 2B, D; Figure 2—figure supplement 1B and D**), and to LAMP1^+^ or cathepsin D^+^ lysosomes (at 24 and 48 h.p.i.) (**Figure 2G-J; Figure 2—figure supplement 1F and H**). Our data, therefore, demonstrated that IREα activity controls Bm16M intracellular replication, likely via regulation of BCV ER trafficking. These findings encouraged us to investigate the molecular mechanisms driving these phenomena.

**Figure 2.**
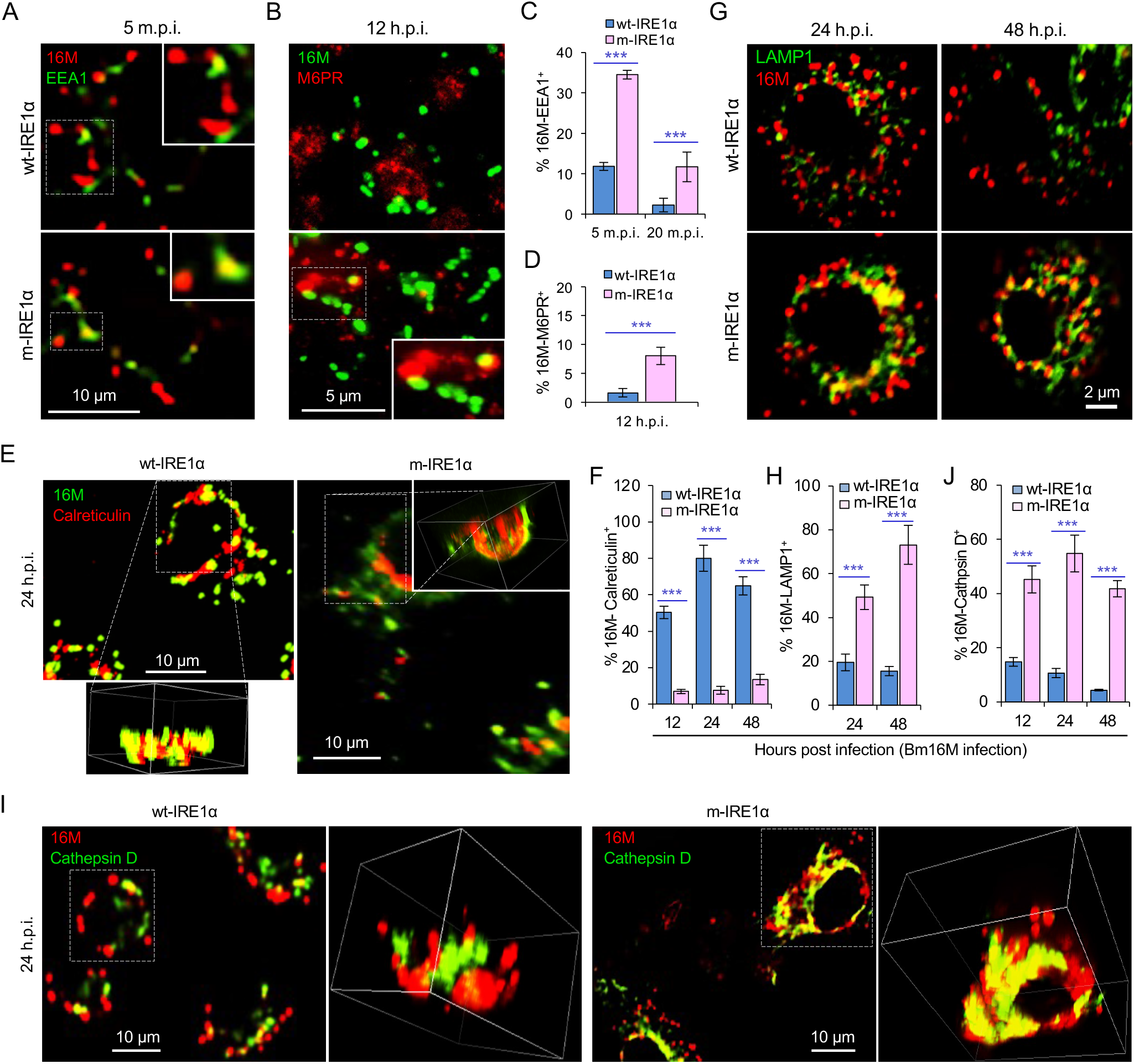
IRE1*α* regulates proper intracellular trafficking and replication of *Brucella* in a XBP1-independent fashion. (A, B) Colocalization analysis of Bm16M with host early endosomes (A) and late endosomes (B) of BMDMs from wt- and m-IRE1α mice at the indicated time points post-infection. m.p.i.: minutes post infection. EEA1: early endosomal antigen 1; M6PR: mannose-6-phosphate receptor. (C, D) Quantification of Bm16M entry into early endosomes (C) or late endosomes (D) of the indicated host BMDMs at the indicated time points post-infection. (E, F) Colocalization analysis of Bm16M and the ER marker calreticulin (E), and quantification of Bm16M-calreticulin^+^ (F) in wt- and m-IRE1α BMDMs at the indicated h.p.i. (G-J) Colocalization of Bm16M and the lysosomal markers LAMP1 (G) or cathepsin D (I), and quantification of Bm16M-LAMP1^+^ (H) or -cathepsin D^+^ (J) in wt- and m-IRE1α BMDMs at the indicated h.p.i.. Images are representative of three independent experiments. Statistical data express as mean ± SEM from three independent experiments. ***, *p* < 0.001. **Figure 2—figure supplement 1**. IRE1α is required for *B. melitensis* properly intracellular trafficking.

### *Brucella* infection down-regulates RIDD genes

We were intrigued with the hypothesis that Bm16M subverts host RIDD activity to promote intracellular parasitism. First, since host UPR/IRE1α RNase activity is induced by *Brucella* effectors secreted by the T4SS of the pathogen (de Jong *et al*., 2013), we tested whether RIDD activity was dependent upon the *Brucella* T4SS. We found that induction of IRE1α RNase activity occurred in a *Brucella* T4SS-dependent fashion (**Figure 3—figure supplement 1A**). Next, we performed RNA-seq analysis to define candidate host genes whose transcripts were subject to RIDD control during *Brucella* infection. Specifically, we used Bm16M to infect triplicate sets of BMDMs as follows: (1) solvent control-treated wt-IRE1α, (2) 4μ8C-treated wt-IRE1α, or (3) solvent control-treated m-IRE1α (**Figure 3—figure supplement 1B**). At 4 or 24 h.p.i., we harvested host mRNA for RNA-seq analysis. Differential expression analysis was then performed to identify genes that were down-regulated following Bm16M infection of wt-IRE1α cells but were unchanged or up-regulated in either infected, drug-treated cells, or infected, m-IRE1α cells. Genes that displayed reduced expression (*p* < 0.05) in response to infection at 4 and/or 24 h.p.i., and also whose infection-dependent reductions in expression were reversed upon treatment with 4μ8C, or in m-IRE1α cells, were defined as candidate “RIDD genes”. This analysis resolved 847 candidate RIDD genes (**Figure 3A-C; Figure 3—figure supplement 1C-F; Figure 3—figure supplement 2; Figure 3—Source Dataset 1**). KEGG pathway and interaction network analyses revealed that most RIDD candidate genes were involved in cellular component organization and biogenesis, RNA metabolism, and oxidative phosphorylation (**Figure 3—figure supplement 2**).

**Figure 3.**
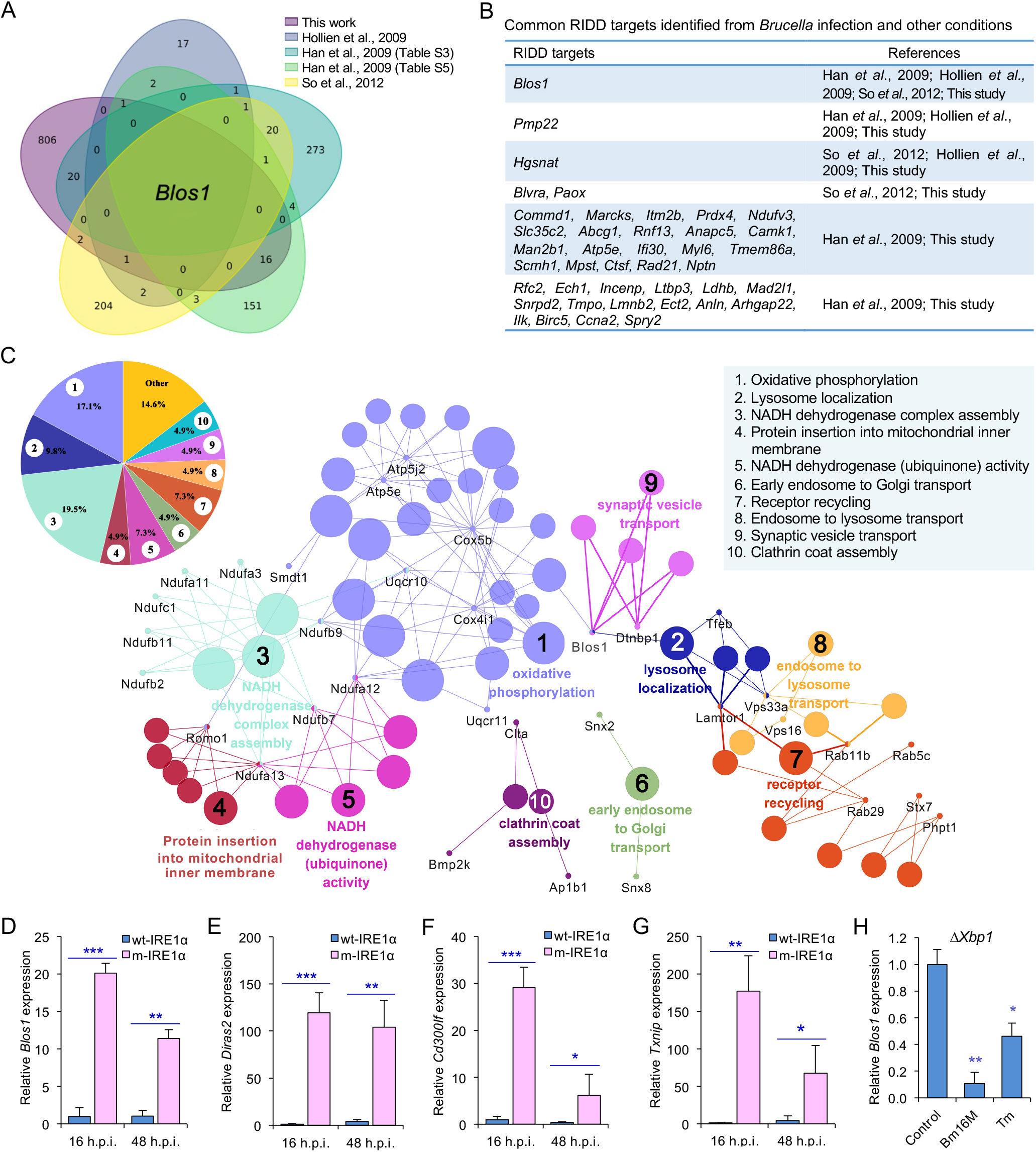
RIDD (regulated IRE1-dependent decay)-BLOS1 axis controls *Brucella* host cell infection. (A) Venn diagram showing numbers of candidate RIDD genes identified in the indicated data sets. (B) Common candidate RIDD genes identified in Bm16M-infected cells and other conditions in the indicated data sets. (C) Interaction network analysis of candidate RIDD genes (*Blos1* associated genes) identified in *Brucella* infected cells and the corresponding enriched KEGG pathways. Different pathways are distinguished by different colors. Interacting genes are shown with the smallest sized of dot with gene names. The upper-left-corner panel: enriched KEGG pathways and interacting candidate RIDD genes (%). (D-G) qRT-PCR validation of RIDD candidate genes *Blos1* (D), *Diras2* (E), *Cd300lf* (F), and *Txnip* (G) identified from RNA-seq analysis. Relative mRNA expression levels in potential RIDD targets from control and m-IRE1α BMDMs infected with Bm16M at 16 and 48 h.p.i were measured by qRT-PCR. (H) qRT-PCR analysis of expression levels of *Blos1* in Δ*Xbp1* BMDMs that were either uninfected (control), infected with Bm16M, or treated with tunicamycin (Tm, an UPR inducer, 5 μg/ml) at 4 hr post infection/treatment. Expression levels of the indicated genes were normalized to GAPDH expression. Statistical data represent the mean ± SEM from three independent experiments. *, **, ***: significance at p < 0.05, 0.01 and 0.001, respectively. **Figure 3—figure supplement 1**. IRE1α activation is *Brucella* Type 4 secretion system (T4SS)-dependent and gene profiling of host cells infected by *Brucella*. **Figure 3—figure supplement 2**. KEGG pathway network analysis of the candidate RIDD genes identified via RNA-seq analysis from host cells infected or uninfected with Bm16M and/or treated or untreated with 4μ8C at 4 and/or 24 h.p.i.. **Figure 3—figure supplement 3**. Validation of RIDD target genes.

We performed several experiments to validate candidate RIDD genes identified in the RNA-seq analysis. First, we used real-time quantitative reverse transcription-PCR (qRT-PCR) to measure the expression levels during infection of several candidate RIDD genes, including *Blos1, Cd300lf, Diras2*, and *Txnip* (**Figure 3A-C; Figure 3—Source Dataset 1**). We found that the expression of these genes was significantly lower in *Brucella* infected wt-IRE1α cells than in m-IRE1α cells (**Figure 3D-G**). Second, we examined whether similar reductions in expression of candidate RIDD genes were observed in host cells infected with *B. abortus* S19 (BaS19, a vaccine strain). We found that BaS19 induced similar phenotypes as Bm16M (**Figure 3—figure supplement 3A-C)**, suggesting that the phenotype was not species-specific. Third, we used qRT-PCR to test the hypothesis that heat-killed bacteria induced similar changes in RIDD gene expression. We found that heat-killed bacteria did not cause a similar effect. Hence, the induction of RIDD activity in host cells required interactions with the viable agent (**Figure 3—figure supplement 3D**) and was also T4SS-dependent (**Figure 3— figure supplement 1A**). Fourth, we compared our list of candidate RIDD genes to genes previously reported to be subject to RIDD control (Bright *et al*., 2015; Han *et al*, 2009; Hollien *et al*., 2009; So *et al*, 2012). This comparison identified 40 genes that were previously shown to be substrates of IRE1α RNase activity and/or displayed expression patterns consistent with RIDD targeting (**Figure 3A-B**). Finally, we found that the expression of the key RIDD gene *Blos1* was also reduced in Δ*Xbp1* BMDMs infected with Bm16M, or when ER stress was induced in these cells (**Figure 3H**). These data suggested that the observed changes in host gene expression patterns were not a consequence of alterations in XBP1 transcription factor activity. Taken together, these data supported the hypotheses that (1) Bm16M infection induces RIDD activity in host cells, and (2) RIDD activity confers enhanced susceptibility to intracellular parasitism by *Brucella*. However, these findings left open the question of the molecular mechanism by which RIDD activity controlled Bm16M replication.

### RIDD activity on *Blos1* controls *Brucella* intracellular parasitism

Our RNA-seq analysis identified *Blos1* as a *Brucella*-induced RIDD gene. However, the mechanisms by which *Blos1* regulates microbial infection are largely unknown. This fact encouraged us to test thehypothesis that *Blos1* plays a central role in regulating Bm16M intracellular parasitism. First, we generated a cell line carrying a non-functional *Blos1* mutant allele (m*Blos1*). Mammalian BLOS1 contains three conserved XAT hexapeptide-repeat motifs that are essential for acetyltransferase activity and may also be a necessary structure-defining feature for acetyl-CoA contact (Scott *et al*, 2018; Wu *et al*, 2021a). Using CRISPR/cas9-mediated gene editing, we mutated the first XAT hexapeptide-repeat motif, which in the wild-type encodes “EALDVH,” and in the mutant encodes “EVVDH or EVDH” (**Figure 4—figure supplement 1A, Table S1**). A cell line containing gene encoding Cas9 and a non-specific gRNA was used as a control of the m*Blos1* mutant line. Second, we generated a RIDD-resistant *Blos1* cell line (henceforth Rr-*Blos1*). In this line, a mutation (from “G” to “U”) was introduced into *Blos1* mRNA stem-loop structure (i.e., the target of IRE1α RNase activity) that rendered resistant cleavage by IRE1α RNase. A *Blos1::Blos1-HA* line that overexpresses *Blos1* (wt-*Blos1*) served as a control of the Rr-*Blos1* cell line (**Figure 4—figure supplement 1B; Table S1**).

**Figure 4.**
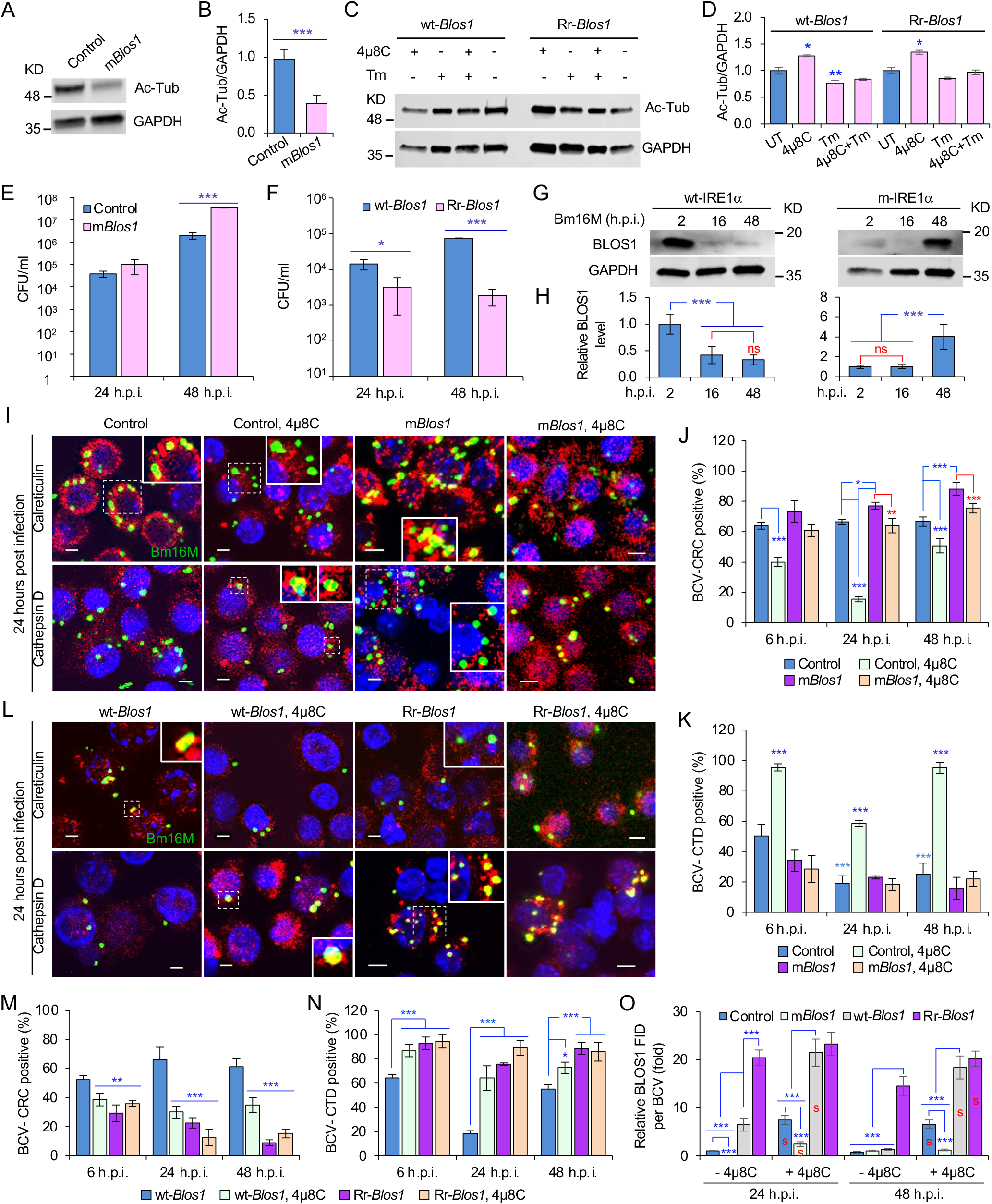
BLOS1 confers host cell susceptibility to *Brucella* infection and controls *Brucella* intracellular trafficking. (A, B) Western blot analysis of α-tubulin acetylation (A) and quantification of α-tubulin acetylation level (B) in control containing Cas9 and a non-specific gRNA and the non-functional *Blos1* mutant (m*Blos1*) in RAW 264.7 Cas9 cells. Ac-Tub: anti-acetylated antibody. (C, D) Western blot analysis of α-tubulin acetylation (C) and quantification of α-tubulin acetylation levels (D) in control (wt-*Blos1*, overexpressing WT *Blos1*) cells and cells that express the RIDD-resistant *Blos1* variant (Rr-*Blos1*) treated or untreated with 4μ8C (50 μM), Tm (5 μg/ml), or both for 4 hr. (E-F) CFU assays for Bm16M infection of RAW264.7 cells in which *Blos1* is non-functional (E), RIDD-resistant (F) at the indicated h.p.i.. (G-H) BLOS1 degradation assay during *Brucella* infection (G) and quantification of the relative BLOS1 expression level (compared to the level of the loading control GAPDH) (H) at the indicated h.p.i.. ns: no significance. (I-K) Colocalization of BCV with calreticulin (CRC) or cathepsin D (CTD) (I) and quantification of BCV-CRC^+^ (J) or BCV-CTD^+^ (K) in control and m*Blos1* cells treated with or without 4μ8C (50 μM) at the indicated h.p.i.. (L-N) Colocalization of BCV with CRC or CTD (L) and quantification of BCV-CRC^+^ (M) or BCV-CTD^+^ (N) in the wt-*Blos1* and Rr-*Blos1* cells treated with or without 4μ8C (50 μM) at the indicated h.p.i.. (O) Quantification of BLOS1 fluorescence integrated density (FID) per BCV in the m*Blos1*, Rr-*Blos1*, or their corresponding control cells treated with or without 4μ8C (50 μM) at the indicated h.p.i.. S: significance (p < 0.01) compared to that without 4μ8C treatment. Host cells were infected with or without Bm16M, and at the indicated h.p.i., the cells were harvested for immunoblotting assays or fixed and subjected to confocal immunofluo-rescence assays. Blots/images are representative of three independent experiments. Statistical data represent the mean ± SEM from three independent experiment. *, p < 0.05; **, p < 0.01; ***, p < 0.001. **Figure 4—figure supplement 1**. Generation of non-functional and overexpression *Blos1* variants. **Figure 4—figure supplement 2**. Cells with BLOS1 deficiency or RIDD resistance differently control lysosome intracellular trafficking. **Figure 4—figure supplement 3**. Host endogenous BLOS1 and the associated proteins are specifically recognized by the indicated home-made or commercial antibodies and Differential interactions of *Brucella* and host BLOS1 during infection.

We characterized the developed cell lines in several ways. First, we noted that α-tubulin acetylation levels had been reported to be controlled, in part, by BLOS1 activity levels (Wu *et al*, 2018). Therefore, we monitored α-tubulin acetylation to assess whether our developed cell lines did, in fact, display alterations in BLOS1 activity. We found that m*Blos1* and Rr-*Blos1* cells displayed reduced levels (**Figure 4A-B**) and maintained relatively higher levels (**Figure 4C-D**), respectively, of acetylated α-tubulin, compared to their corresponding controls. These data supported the hypothesis that these cells had the expected levels of BLOS1 activity. Second, we tested the replication of the pathogen in different *Blos1* cell lines. We found that m*Blos1* cells exhibited increased susceptibility to Bm16M infection (**Figure 4E**), whereas Rr-*Blos1* cells or wild-type controls treated with 4μ8C supported dramatically reduced intracellular bacterial growth (**Figures 1M; 4F; Figure 4—figure supplement 1C**). Finally, we monitored the expression levels of BLOS1 protein during a 48 hr time course of infection. We found that BLOS1 expression was reduced at 16 h.p.i., and continuously decreased during Bm16M infection in wt-IRE1α control cells; however, in m-IRE1α BMDMs, BLOS1 expression was relatively stable or increased (at 48 h.p.i.) (**Figure 4G-H**). Similar results were observed in 4μ8C treated or untreated m*Blos1*, Rr-*Blos1*, and control cells infected with BaS2308 (**Figure 4—figure supplement 1D-E**). These data demonstrate that low or high BLOS1 expression levels promote or impair *Brucella* infection, respectively.

### BLOS1 regulates *Brucella* intracellular trafficking

The mechanism by which BLOS1 regulates *Brucella* infection was unknown. However, the observed subcellular trafficking defect of the pathogen in host cells harboring mutant or deficient variants of IREα (**Figure 2; Figure 2—figure supplement 1**) suggested that BLOS1 may control the intracellular parasitism of the pathogen by regulating its subcellular trafficking. To illuminate this aspect, we treated m*Blos1*, Rr-*Blos1*, and the corresponding control cell lines with tunicamycin (Tm, an UPR inducer) or 4μ8C, or infected them with Bm16M. We then assessed the trafficking of the pathogen in these cells using CIM. We found that low levels of BLOS1 or non-functional BLOS1 in uninfected or infected cells were associated with the accumulation of late endosome/lysosome (LE/Lys) membranes in the vicinity of nuclei, reduced colocalization of latex beads with cathepsin D, and increased perinuclear LC3b index or autophagic activity near nuclei, in both control and Tm-treated conditions (**Figure 4—figure supplement 2A, C, E, G**). In these studies, the LC3b index was defined as: (Total number of identified cells with the ratio of the mean LC3b intensity in the cytoplasm to that in the nucleus <1)/(Total number of the analyzed cells).

In contrast, overexpression of BLOS1 (the wild-type cells expressing wt-*Blos1* or Rr-*Blos1*) reduced LE/Lys perinuclear accumulation, increased the localization of latex beads in cathepsin D^+^ compartments, and reduced perinuclear autophagic activity (**Figure 4—figure supplement 2B, D, F, H**). Although significant inhibition of BCV trafficking to lysosomes in the m*Blos1* cells was not observed at 24 and 48 h.p.i., the m*Blos1* cells supported enhanced BCV trafficking to ER compartments during bacterial infection, compared to that in the wild-type control cells, or 4μ8C-treated m*Blos1* cells (**Figure 4I-K**). In contrast, Rr-*Blos1* cells displayed reduced BCV trafficking to ER compartments, but instead promoted BCVs trafficking to lysosomes during Bm16M infection, compared to controls (**Figure 4L-N**).

To test the hypothesis that Bm16M infection alters the dynamics of associations between BLOS1 and BCVs, we used CIM approaches to localize these elements during a time course of infection after confirmation of the specificities of antibodies used in the work (see below) (**Figure 4—figure supplement 3A-D**). We found higher levels of BLOS1 colocalization with BCVs in 4μ8C-treated control or m*Blos1* cells than their corresponding untreated cells (**Figure 4O**); moreover, at 24 h.p.i., lower levels of BLOS1^+^ BCVs were observed in m*Blos1* cells compared to wild-type controls (**Figure 4—figure supplement 3E-F**). Rr-*Blos1* and/or 4μ8C inhibition of *Blos1* degradation in host cells (i.e., Rr-*Blos1*, 4μ8C-treated wt-*Blos1* control or Rr-*Blos1* cells) significantly promoted BLOS1^+^ BCVs compared to the wt-*Blos1* control (**Figure 4O,Figure 4—figure supplement 3E-F**). These findings demonstrated that non-functional BLOS1 promoted the trafficking of the pathogen from LE/Lys membranes to the ER. However, Rr-*Blos1* cells (or 4μ8C treated cells) promoted the trafficking and degradation of the pathogen in lysosomes.

### Disassembly of BORC promotes BCV trafficking to and accumulation in the ER

To test whether BORC-related lysosome trafficking components mediate BCV trafficking during infection, we analyzed the dynamics of the interaction of BCVs with LAMTOR1 (a central component of mTORC1), the small GTPase ARL8b, and kinesin KIF1b and KIF5b proteins during infection. During bacterial intracellular trafficking and replication, colocalization of BCVs with both LAMP1 and LAMTOR1, LAMP1, or LAMTOR1 decreased in control and m*Blos1* cells, whereas these interactions were observed at higher levels in cells harboring Rr-*Blos1* variants (**Figure 5A-B, E** and **G; Figure 5—figure supplement 1A-B**). Similarly, recruitment of ARL8b to BCVs and/or LAMP1 was reduced in control and m*Blos1* cells, which impaired their kinesin-dependent movement toward the cell periphery; however, these interactions were maintained in Rr-*Blos1* cells (**Figure 5C-D, F** and **H; Figure 5—figure supplement 1C-D**). BCV interactions with KIF1b^+^ or KIF5b^+^, which preferentially drive lysosomes on peripheral tracks or perinuclear/ER tracks (Guardia *et al*., 2016), decreased (**Figure 6A-B, E** and **G; Figure 6—figure supplement 1A-B**) or increased (**Figure 6C-D,F** and **H; Figure 6—figure supplement 1C-D**), respectively, in control, m*Blos1*, and wt-*Blos1* cells; however, the opposite phenomena were observed in Rr-*Blos1* cells (**Figure 6; Figure 6—figure supplement 1**). These findings suggest that BORC-related lysosome trafficking components may regulate BCV perinuclear trafficking, fusion with ER membranes and subsequent bacterial replication.

**Figure 5.**
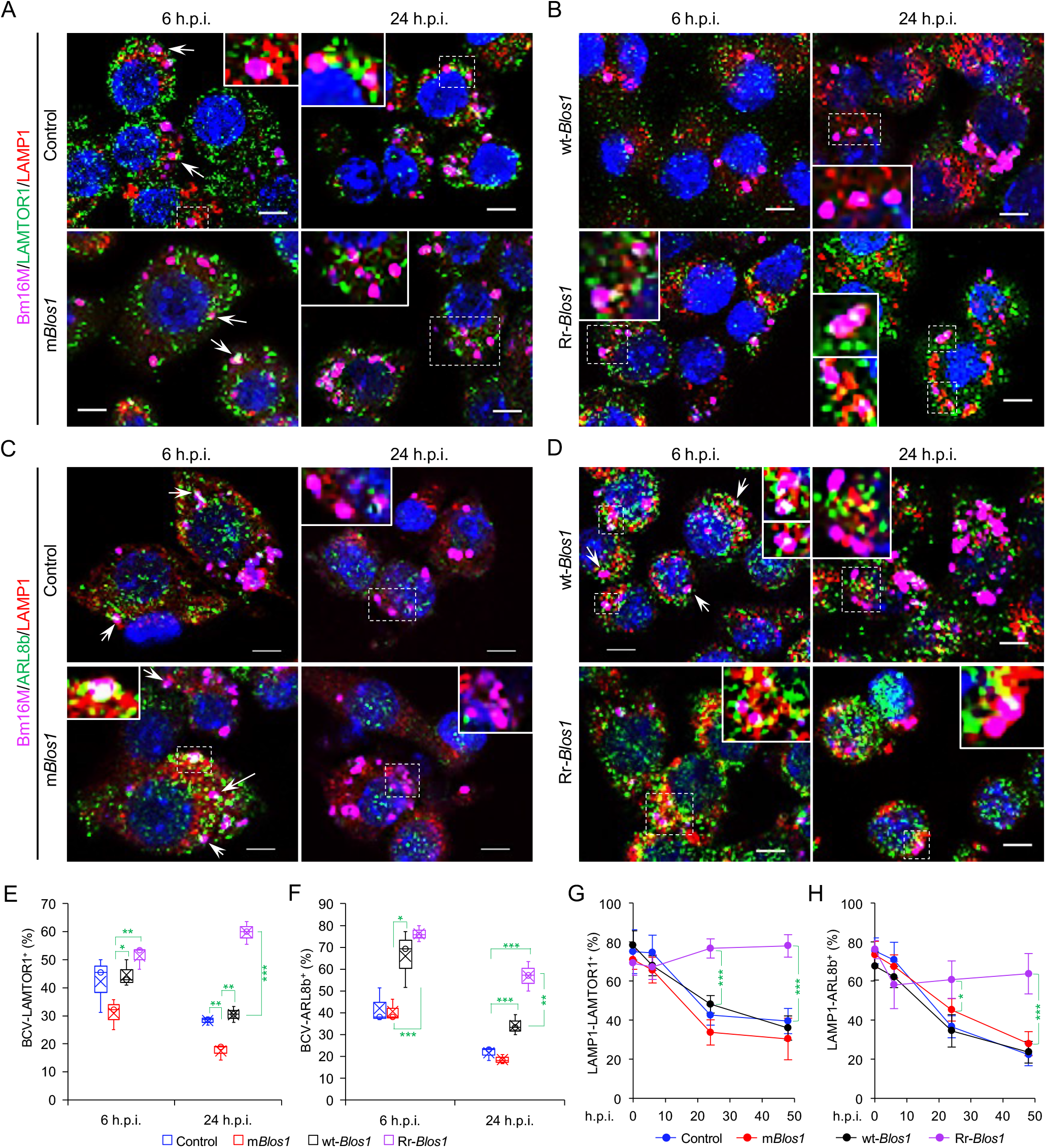
*Brucella* infection dissociates host BORC-related lysosome trafficking factor LAMTOR1 and ARL8b from lysosomes. (A, B) Colocalization of LAMTOR1 with BCVs or LAMP1 in the infected control and m*Blos1* (A), or in wt-*Blos1* and Rr-*Blos1* (B) cells at the indicated h.p.i.. (C, D) Colocalization of ARL8b with BCVs or LAMP1 in the infected control and m*Blos1* (C), or in wt-*Blos1* and Rr-*Blos1* (D) cells at the indicated h.p.i.. Arrows: colocalization of BCVs with the indicated proteins. Insets: magnification of the selected areas (within windows with dash white lines). Bars: 5 μm. (E-F) Quantification of BCV-LAMTOR1^+^ (E) and BCV-ARL8b^+^ (F) in Bm16M infected cells at the indicated h.p.i. showing in (A-B) and (C-D), respectively. (G-H) Dynamics of LAMP1-LAMTOR1^+^ (E) or LAMP1-ARL8b^+^ (H) in a time course (48 hr) of Bm16M infection at the indicated h.p.i.. Host cells were infected with or without Bm16M, and at the indicated h.p.i., the cells were fixed and performed confocal immunofluorescence assays. Images are representative of three independent experiments. Statistical data expressed as mean ± SEM from three independent experiments. *, p < 0.05; **, p < 0.01; ***, p < 0.001. **Figure 5—figure supplement 1**. Association of the indicated BORC-related lysosome trafficking components in the indicated uninfected-host cells.

**Figure 6.**
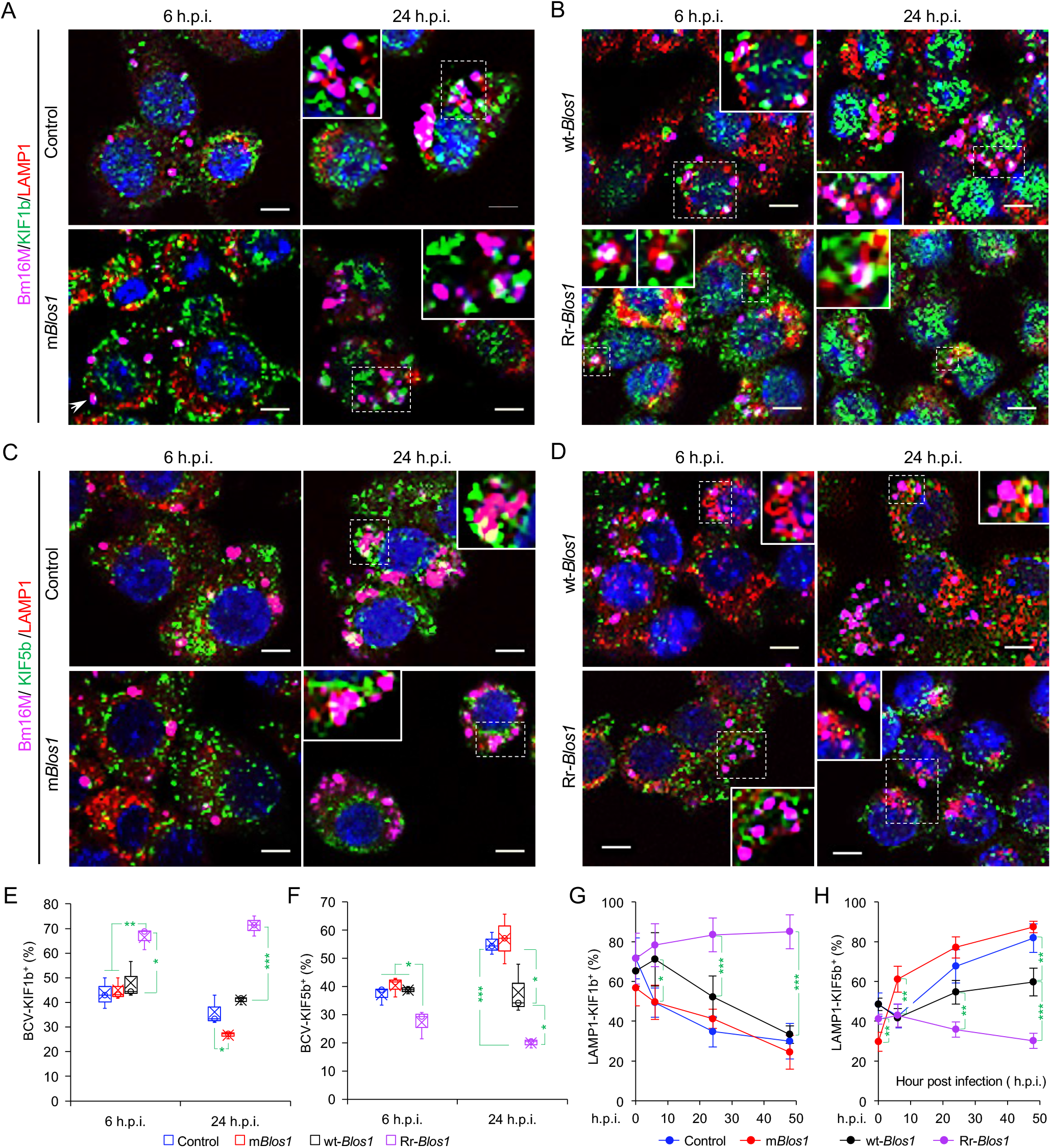
*Brucella* infection dissociates BORC-related lysosome trafficking factor KIF1b but recruits KIF5b. (A, B) Colocalization of KIF1b with BCVs or LAMP1 in the infected control and m*Blos1* (A), or in wt-*Blos1* and Rr-*Blos1* (B) cells at the indicated h.p.i.. (C, D) Colocalization of KIF5b with BCVs or LAMP1 in the infected control and m*Blos1* (C), or in wt-*Blos1* and Rr-*Blos1* (D) cells at the indicated h.p.i.. Arrows: colocalization of BCVs with the indicated proteins. Insets: magnification of the selected areas (within windows with dash white lines). Bars: 5 μm. (E-F) Quantification of BCV-KIF1b^+^ (E) and BCV-KIF5b^+^ (F) in Bm16M infected cells at the indicated h.p.i. showing in (A-B) and (C-D), respectively. (G-H) Dynamics of LAMP1-KIF1b^+^ (G) or LAMP1-KIF5b^+^ (H) in a time course (48 hr) of Bm16M infection at the indicated h.p.i.. Host cells were infected with or without Bm16M, and at the indicated h.p.i., the cells were fixed and subjected to confocal immunofluorescence assays. Images are representative of three independent experiments. Statistical data represent means ± SEM from three independent experiments. *, p < 0.05; **, p < 0.01; ***, p < 0.001. **Figure 6—figure supplement 1**. Association of the BORC-related lysosome trafficking components KIF1b, KIF5b, and LAMP1 in the indicated uninfected-host cells.

BORC, a protein complex that contains three components (i.e., BLOS1, BLOS2, SNAPIN) shared with the BLOC-1 complex and five other proteins KXD1, C17orf59 (Lyspersin), LOH12CR1(Myrlysin), C10orf32 (Diaskedin), and MEF2BNB, plays a critical role in the regulation of lysosome positioning (Pu *et al*., 2015). In HeLa cells, interference with BORC triggers LE/Lys trafficking to the cell center via dynein, resulting in a characteristic clustering of LE/Lys in perinuclear regions (Pu *et al*., 2015). We hypothesized that degradation of *Blos1* mRNA by IRE1α during *Brucella* infection interferes with BORC assembly, resulting in the alteration of recruitment or disassociation of BORC-related trafficking components, and increased LAMP1^+^-BCV perinuclear trafficking and fusion with the ER and/or macroautophagosome membranes. To test this hypothesis, we performed protein co-immunoprecipitation (Co-IP) assays to measure the association of BORC components with each other in *Brucella* infected or uninfected host cells. We found that in uninfected cells, BLOS1 interacted with protein components of BLOC-1 (PALLIDIN), BORC (KXD1), and both BLOC-1 and BORC (BLOS2, SNAPIN) (**Figure 7A and B**). Moreover, under this condition, BORC components localized with peripheral or cytosolic LE/Lys membranes (**Figure 5—figure supplement 1; Figure 6—figure supplement 1**). However, in *Brucella* infected cells, where *Blos1* mRNA was degraded and BLOS1 protein depletion was observed (**Figures 3D, H; 4G-H; Figure 3—figure supplement 3A; Figure 4—figure supplement 1D-E**), physical interaction between BLOS1 and BORC component SNAPIN was reduced in control cells and difficult to detect in the m*Blos1* variants during intracellular trafficking and replication of the pathogen (48 h.p.i.) (**Figure 7C-D**). In fact, interactions between LYSPERSIN and KXD1 in control cells were also only detected at early time points (2 h.p.i.), but not at later time points corresponding to intervals when bacterial intracellular trafficking and replication were expected to occur; these interactions were also hardly detected in infected cells expressing m*Blos1* variants at these time points (**Figure 7C and E**). The reduced interactions between BLOS1 and BORC component SNAPIN as well as LYSPERSIN and KXD1 may result from the disassembly of BORC when BLOS1 is degraded during *Brucella* infection (**Figure 7C-E**). In *Brucella* infected *Blos1* overexpressing (wt-*Blos1*) cells, substantial reductions in the interactions were also observed (**Figure 7F-G**). In the infected Rr-*Blos1* cells, the interaction was maintained at a relatively higher level (**Figure 7F-G**), suggesting that the BORC complex remains assembled. The integrity or disassociation of BORC was consistent with the interactions between BORC-related trafficking components and with the colocalization dynamics of BCVs with BLOS1, mTORC1/LAMP1, LAMP1/ARL8b, LAMP1/KIF1b or with LAMP1/KIF5b. These findings were also consistent with BCV peripheral or perinuclear/ER trafficking and accumulation (**Figure 5-6; Figure 4—figure supplement 3E-F**). The results collectively suggested that the degradation of *Blos1* mRNA during pathogen infection resulted in the disassembly of BORC, which promoted BCV trafficking to and accumulation in the vicinities of nuclei and likely facilitated the fusion of BCVs with ER membranes in which bacteria replicated.

**Figure 7.**
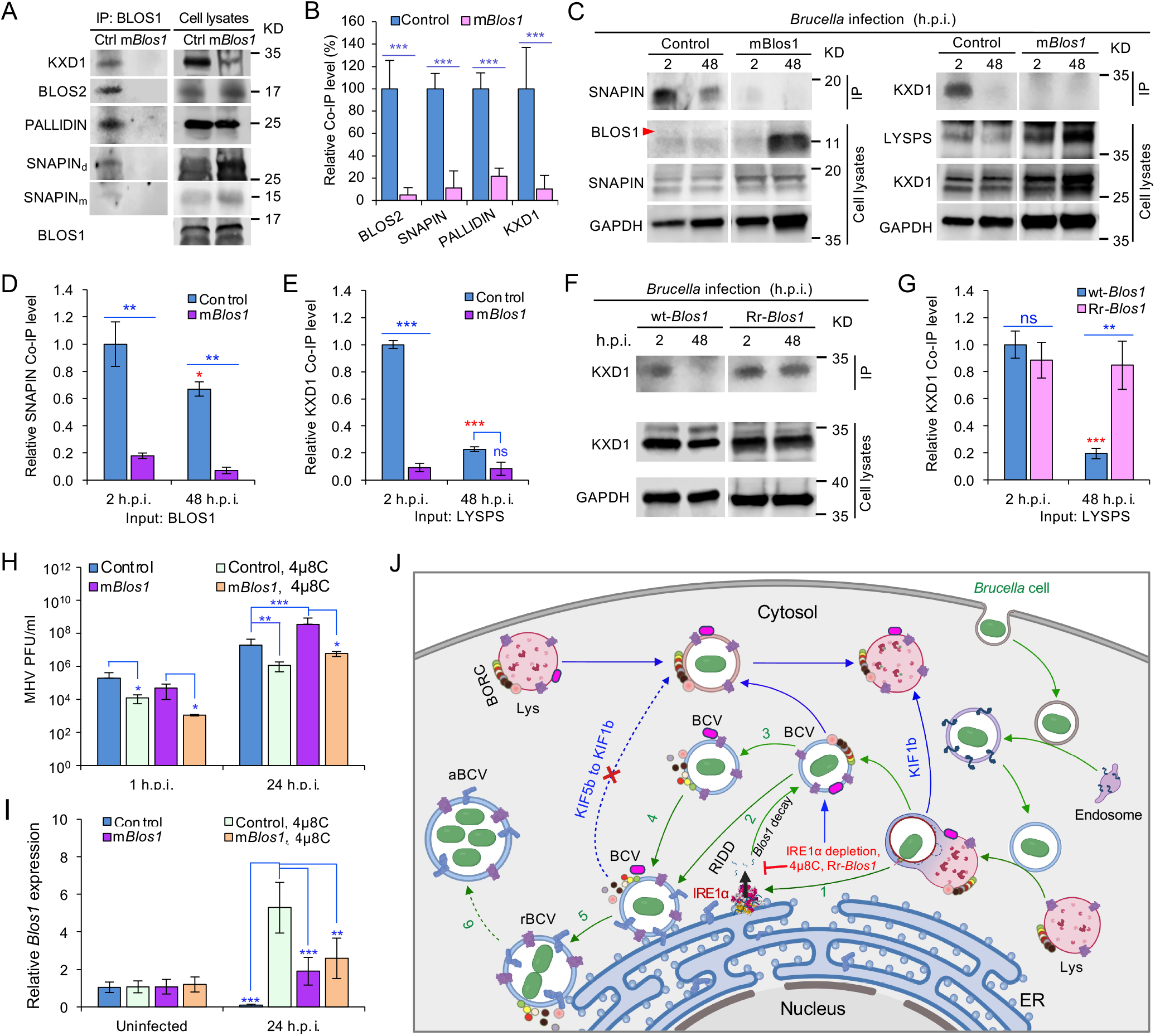
*Brucella* infection dissociates host BORC and degradation of *Blos1* mRNA supports coronavirus intracellular replication. (A) Co-immunoprecipitation (Co-IP) analysis of the interactions of BLOS1 with other proteins that form the BLOC-1 (BLOS1, BLOS2, SNAPIN, and PALLIDIN) or BORC (BLOS1, BLOS2, SNAPIN, and KXD1) complex. The non-functional m*Blos1* and control cells were cultured overnight before being subjected to Co-IP assays with BLOS1 as an input. SNAPIN_m_: monomer SNAPIN; SNAPIN_d_: dimer SNAPIN. (B) Quantification of the indicated pulled-down protein levels from overnight-cultured control and m*Blos1* cell lysates using BLOS1 as an input. (C) Co-IP assays for *Brucella*-infected control and m*Blos1* cells at the indicated h.p.i. using BLOS1 (leaf panel) or LYSPERSIN (LYSPS, right panel) as an input. Red arrow: BLOS1. (D-E) Quantification of the indicated pulled-down protein levels of SNAPIN (D) or KXD1 (E) from *Brucella*-infected control and m*Blos1* cell lysates using BLOS1 or LYSPS as an input. (F) Co-IP assays for *Brucella*-infected wt- or Rr-*Blos1* cells at the indicated h.p.i. using LYSPERSIN as an input. (G) Quantification of the indicated pulled-down protein levels from *Brucella*-infected cell lysates of wt- or Rr-*Blos1*. (H) PFU (plaque-forming units) assay of coronavirus MHV infection of the m*Blos1* and control cells treated or untreated with 4μ8C (50 μM) at the indicated h.p.i.. (I) *Blos1* mRNA expression assay of the indicated host cells infected with MHV via qRT-PCR. (J) A proposed model describing how *Brucella* subverts the host RIDD-BLOS1 pathway to support intracellular parasitism by disrupting BORC-directed lysosomal trafficking. Green arrows: BCV trafficking to the ER compartment and replication. Blue arrows: BCV trafficking to peripheral lysosome and lysosomal degradation. Host cells were infected with or without *Brucella* or MHV, and at the indicated h.p.i., the infected or uninfected host cells were harvested for Co-IP and immunoblotting assays, or qRT-PCR assays. Blots are representative of three independent experiments. Statistical data represent the mean ± SEM from three independent experiment. *, p < 0.05; **, p < 0.01; ***, p < 0.001. Red asterisks: Compared to the same *Brucella* infected cells at 2 h.p.i..

### Host RIDD activity on BLOS1 promotes coronavirus intracellular replication

In light of the global COVID-19 pandemic, we tested whether RIDD-controlled BLOS1 activity is a target for subversion by coronaviruses. We infected control or host cells harboring alterations in this pathway with mouse hepatitis virus [MHV, a positive-strand RNA virus classified as a member of the Betacoronavirus genus (CoV)]. Notably, previous studies have shown that MHV infection induces host cell UPR and activates IRE1α RNase and *Xbp1* splicing (Bechill *et al*, 2008), thereby suggesting the hypothesis that MHV infection of host cells activates RIDD activity. To test this hypothesis, m*Blos1* or control host cells were untreated or treated with 4μ8C. Next, these cells were infected with MHV for 24 hr. Virus plaque-forming units (PFU) and host *Blos1* expression were then measured. We found that viral PFUs were reduced in 4μ8C-treated cells. However, significantly increased PFU in m*Blos1* cells at 24 h.p.i. compared to controls was observed (**Figure 7H**). Expression levels of *Blos1* mRNA were dramatically reduced during infection (**Figure 7I**). Collectively, these findings suggested that coronavirus MHV, like *Brucella*, subverts the host RIDD pathway to promote intracellular infection.

## Discussion

RIDD, a fundamental component of UPR in eukaryotic cells, cleaves a cohort of mRNAs encoding polypeptides that influence ER stress, thereby supporting the maintenance of ER homeostasis. In this report, we found that *Brucella* infection subverts UPR, in general (Pandey *et al*., 2018; Qin *et al*., 2008; Smith *et al*., 2013; Taguchi *et al*., 2015), and RIDD activity on *Blos1*, in particular, to promote intracellular parasitism. BLOS1, encoded by RIDD gene *Blos1*, is a shared subunit of both BLOC-1 and BORC complexes (Pu *et al*., 2015). Mutation or a reduction in BLOS1 expression affects both BLOC-1 and BORC (**Figure 7A-H**). The BLOC-1 complex is mainly involved in endosomal maturation and endosome-lysosome trafficking and fusion (John Peter *et al*, 2013; Pu *et al*., 2015; Scott *et al*., 2018). Our work does not rule out the possibility that the disassociation of BLOC-1 also affects *Brucella* intracellular parasitism, especially in the early stages of cellular infection.

Our findings support a stepwise working model by which *Brucella* subverts the host RIDD pathway to facilitate intracellular parasitism (**Figure 7J**). First, *Brucella* infection induces UPR in host cells, a process associated with activation of IRE1α kinase (Pandey *et al*., 2018; Taguchi *et al*., 2015) and RNase activities (Smith et al., 2013; this work). Second, degradation of the RIDD target *Blos1* by IRE1α RNase activity results in depletion of BLOS1 proteins and reduced association with BORC components (**Figures 3D; 4G-H; Fig 7A-H**). Third, these events drive the trafficking of BCVs to the ER and their perinuclear accumulation, mitigate further fusion of BCVs with cytosolic lysosomes, and limit BCV trafficking to LE/Lys in peripheral regions where these organelles possess enhanced degradative functions (**Figures 5-6; Figure 4—figure supplement 3E-F**). Finally, the accumulation of BCVs decorated with ER proteins increases due to the fusion of BCVs with ER membranes and/or with noncanonical macrophagosomes (Pandey *et al*., 2018; Starr *et al*., 2012; Taguchi *et al*., 2015). These final events support the intracellular replication and cell-to-cell movement of the pathogen.

Several lines of evidence support the proposed mechanism. First, *Brucella* infection activates IRE1α RNase activity as evidenced by *Xbp1* mRNA splicing (**Figure 3—figure supplement 1A**). However, the intracellular replication of the pathogen is not impaired in *Xbp1* KO cells and mice (**Figure 1N-P**). These findings support the hypothesis that IRE1α RNase activity is required for *Brucella* infection in an IRE1α-XBP1 independent fashion. Second, in addition to splicing *Xbp1*, IRE1α cleaves other mRNAs, resulting in their RIDD-mediated decay (Bae *et al*., 2019). We identified several mRNAs, including *Blos1* (**Figure 3A-C**), that contain predicted stem-loop structures that were inferred to be targets of IRE1α RNase activity (Moore & Hollien, 2015). The expression of these mRNAs was downregulated in response to *Brucella* infection. Host cells harboring non-functional *Blos1* mutants were highly susceptible to pathogen infection, whereas cells that express a RIDD-resistant *Blos1* variant were resistant to *Brucella* infection (**Figure 4E-F**).

Third, lysosome positioning regulated by BORC is a critical determinant of its functions. BORC associates peripherally with lysosomal membranes, where it recruits the small GTPase ARL8b to lysosomes. BORC and ARL8b promote lysosome movement by coupling to kinesin-1 (KIF5b) or kinesin-3 (KIF1b), which preferentially moves lysosomes on perinuclear tracks enriched in acetylated α-tubulin or on peripheral tracks enriched in tyrosinated α-tubulin, respectively (Guardia *et al*., 2016; Pu *et al*., 2015). Interference with BORC or other components of this pathway drives lysosome trafficking to the cell center via dynein. Thus, cells lacking BORC display a perinuclear clustering of lysosomes (Pu *et al*., 2015). Ragulator (a GEF for the Rag GTPases that signal amino acid levels to mTORC1) directly interacts with and inhibits BORC functions (Pu *et al*., 2017). Building upon these observations, we show that *Brucella* infection results in *Blos1* degradation and disassembly of BORC; moreover, during *Brucella* intracellular trafficking and replication, colocalization of mTORC1, ARL8b, and KIF1b with BCVs or lysosomes was reduced in control cells, and in cells expressing non-functional variants of BLOS1; however, KIF5b localization with BCVs or lysosomes was increased or in a higher level in these cells (**Figure 5,Figure 5—figure supplement 1; Figure 6,Figure 6—figure supplement 1**). Colocalization of the BORC-related lysosome trafficking factors (i.e., ARL8b, KIF1b, and mTORC1) with BCVs or lysosomes in cells expressing Rr-*Blos1* variants were maintained at a relatively higher level (**Figures 5-6**). These findings demonstrate that blocking BORC function via the disassembly of the BORC complex through depletion of *Blos1* by *Brucella* infection drives BCVs towards the perinuclear region and ER accumulation, which likely facilitates the fusion of BCVs with the ER, thereby supporting intracellular parasitism.

Finally, RIDD-mediated *Blos1* degradation may promote BCV fusion with autophagosomes. Nutrient-starved cells display perinuclear clustering of lysosomes, which influences autophagosome formation and autophagosome-lysosome fusion rates (Korolchuk *et al*, 2011). Lysosome perinuclear clustering during starvation, ER stress induced by accumulation of misfolded proteins, drug treatments, and pathogen infection can disrupt metabolic homeostasis, thereby necessitating the induction of cell biological processes that return the cell to equilibrium. Macroautophagy and degradation of sequestered cytosolic materials by fusion of autophagosomes/macrophagosomes with lysosomes can promote the re-establishment of homeostasis (Bae *et al*., 2019; Korolchuk *et al*., 2011; Pu *et al*., 2017). Degradation of *Blos1* mRNA by IRE1α leads to the perinuclear accumulation of LE/Lys in response to ER stress in mouse cells. Overriding *Blos1* degradation results in ER stress sensitivity and the aggregation of ubiquitinated proteins. The independent perinuclear-trafficking and LE-associated endocytic transport promote the efficient degradation of these protein aggregates. Therefore, *Blos1* regulation via RIDD facilitates LE-mediated autophagy of protein aggregates, thereby promoting cell survival during stress (Bae *et al*., 2019). Hepatocytes from *Blos1* liver-specific knockout (LKO) cells accumulate autolysosomes and lysosomes. In LKO hepatocytes, the initiation or extension of lysosomal tubules is abolished, which impairs autophagic lysosome reformation and results in the accumulation of enlarged autolysosomes (Wu *et al*, 2021b). *Blos1* degradation by the RIDD pathway promotes BCV perinuclear or ER-region clustering, and may also avoid the peripheral movement of BCVs away from the ER region as a consequence of reduced α-tubulin acetylation. These processes may facilitate BCV fusion with ER membranes or (macro)phagosomes, promote the enlargement of aBCVs and further bacterial replication, and ultimately relieve *Brucella* induced ER stress (Pandey *et al*., 2018; Qin *et al*., 2008; Starr *et al*., 2012; Taguchi *et al*., 2015).

In addition to *Brucella*, the betacoronavirus MHV also subverts the RIDD-BLOS1 axis to promote intracellular replication (**Figure 7H-I**), thereby indicating that RIDD control of BLOS1 activity is not pathogen-specific. How the host RIDD-*Blos1* axis regulates interactions between host cells and coronaviruses merits further investigation. However, additional possibilities for regulatory control can be envisioned. First, coronaviruses utilize many proteins such as nsp1 to inhibit host protein synthesis in the first 6 hr of infection (Nakagawa & Makino, 2021). Second, BLOS1 contains a potential coronavirus 3C-like protease cleavage site, LQ^SAPS, near its C-terminus, thereby rendering it potentially susceptible to direct subversion by coronaviral pathogens. Finally, coronaviruses have evolved to subvert host interferon defenses (Thoms *et al*, 2020), which may contribute to immune evasion. Future work will be directed toward examining these possibilities and the roles and mechanisms by which the RIDD-*Blos1* axis controls these and other host-pathogen interactions.

## Materials and Methods

### Bacterial strains, cell culture, *Brucella* infection and antibiotic protection assays

*Brucella melitensis* strain 16M (WT), and *B. abortus* strain 2308 (WT), and *B. abortus* vaccine strain S19 and other bacterial strains were used in this work. Bacteria were grown in tryptic soy broth (TSB) or on tryptic soy agar (TSA, Difco™) plates, supplemented with either kanamycin (Km, 50 μg/ml) or chloramphenicol (Cm, 25 μg/ml) when required. For infection, 4 ml of TSB was inoculated with a loop of bacteria taken from a single colony grown on a freshly streaked TSA plate. Cultures were then grown with shaking at 37°C overnight, or until OD_600_≈3.0.

Mammalian host cells including murine macrophages RAW264.7 and its derived non-functional and Rr-Blos1 variants and corresponding control cells, BMDMs, J774.A1 cells, MEFs and were routinely cultured at 37°C in a 5% CO_2_ atmosphere in Dulbecco’s Modified Eagle’s Medium (DMEM) supplemented with 10% fetal bovine serum (FBS). Murine osteoblasts MC3T3-E1 and its derived Rr-Blos1 variant and corresponding control cells (generously provided by the Hollien Lab) were routinely cultured at 37°C in a 5% CO_2_ atmosphere in alpha minimum essential media (MEMα) with nucleosides, L-glutamine, and no ascorbic acids, supplemented with 10% FBS. Murine fibroblasts L2 cells were routinely cultured at 37°C in a 5% CO_2_ atmosphere in F12 medium supplemented with 10% fetal calf serum (FCS). For BMDMs, the above-mentioned DMEM with 20% L929 cell supernatant, 10% FBS, and antibiotics was used. Cells were seeded in 24-well or 96-well plates and cultured overnight before infection. For antibiotic protection assays, 1.25×10^5^ (BMDMs) or 2.5×10^5^ (RAW264.7) host cells were seeded in each well; for fluorescence microscopy assays, 1×10^4^ or 5×10^4^ cells were seeded in 96-well plates or on 12-mm glass coverslips (Fisherbrand) placed on the bottom of 24-well microtiter plates respectively; for host RNA analysis, 1×10^5^ host cells were seeded in each well of 24-well plates before infection. Host cells were infected with *Brucella* at an MOI of 100, unless otherwise indicated. Infected cells were then centrifugated for 5 min (200 × g) and incubated at 37°C. Thirty minutes to 1 hr post-infection, culture media was removed, and the cells were rinsed with 1 × phosphate buffered saline (PBS, pH 7.4). Fresh media supplemented with 50 μg/ml gentamicin was then added for 1 hr to kill extracellular bacteria. Infected cells were continuously incubated in the antibiotic. At the indicated time points post infection, viable bacteria in infected cells were analyzed using the antibiotic protection assay or the immunofluorescence microscopy assay as previously described (Pandey *et al*., 2018; Qin *et al*., 2008).

### Viral propagation, infection, and plaque assay

Wild type MHV-A59 was propagated in L2 cells in F12 media with 2% FCS. Host cells (RAW 264.7) were infected with MHV-A59 in triplicate at a MOI of 1. Infected cells were incubated at room temperature with gentle rocking for 1 hour. Afterwards, culture media was removed, and the cells were rinsed with 1× PBS (pH 7.4). Fresh media supplemented with 2% FBS was added. Infected host cells were incubated at 37°C. At the indicated time points post infection, viral supernatants were collected and then titrated by plaque assay on L2 cells at 33°C.

### Generation, genotyping and characterization of Lysm-IRE1*α*^-/-^ mice

Animal research was conducted under the auspices of approval by the Texas A & M University Institutional Animal Care and Use Committee in an Association for Assessment and Accreditation of Laboratory Animal Care International Accredited Animal Facility. To investigate the roles of IRE1α in controlling Bm16M intracellular parasitism, LysM-IRE1α^-/-^-[IRE1α Conditional KO (CKO) mice were generated by breeding mice, in which exon 20-21 was floxed (Iwawaki *et al*., 2009) with lysozyme M (LysM) transgenic mice (Jackson Laboratories, Inc.). In the resultant animals, exon 20-21 of the IRE1α gene was specifically deleted in myeloid cells, including macrophages, monocytes, and neutrophils. The CKO mice were genotyped using genomic DNA from tail vain to show the presence of cre alleles (Iwawaki *et al*., 2009). Western blot analysis using anti-IRE1α antibodies (Novus Biologicals) and *Xbp1* splicing were performed on BMDMs from CKO and control mice to validate the absence of full-length IRE1α in CKO mice.

### BMDM harvest and cultivation

BMDMs collected from the femurs of IRE1α CKO and control mice were cultivated in L929-cell conditioned media [DMEM medium containing 20% L929 cell supernatant, 10% (v/v) FBS, penicillin (100 U/ml) and streptomycin (100 U/ml)]. After 3 days of culture, non-adherent precursors were washed away, and the retained cells were propagated in fresh L929-cell conditioned media for another 4 days. BMDMs were split in 24-well plates (2.5×10^5^ cells/well) in L929-cell conditioned media and cultured at 37°C with 5% CO2 overnight before use.

### Whole animal infections with *Brucella* and tissue analysis

Mice from CKO and littermate control groups were intranasally infected with *B. melitensis* and *B. abortus* (Bm16M and BaS2038, respectively) with a dose of 1×10^6^ CFU. At 7 and 14 dpi, infected mice were euthanized, and the bacterial burden was assessed in spleen and liver. A portion of the tissue was fixed, and paraffin embedded for histopathological examination following H&E staining. To assess Bm16M tissue burden, spleen or liver tissues were homogenized and subjected to a serial dilution. Finally, the diluted tissue homogenates (200 μl) were plated on TSA solid plates and CFUs were determined at 48 to 60 hr post incubation at 37°C in 5% CO_2_.

### Latex bead phagocytosis assays

Phagocytosis assays for testing the phagocytic uptake and route of a substrate in the non-functional and RIDD resistant Blos1 variants in RAW264.7 murine macrophages were performed using the Phagocytosis Assay Kit (IgG FITC) (Cayman Chemical, USA) according to the manufacturer’s instructions.

### RNA extraction and qRT-PCR analysis

RNA was extracted from host cells per instructions in the RNeasy Mini Kit (74134 Qiagen). Complementary DNA was amplified from mRNA using the High-Capacity cDNA Reverse Transcription Kit (4368813 Applied Biosystems) per manufacturer’s guidelines. For qRT-PCR of the macrophage infections of MHV strain A59, BaS19, Bm16M, or Ba2308 1/5 dilution of each cDNA was added into nuclease free water in a respective well in a 96-well plate. SYBR green (50 - 90 μl) with primers (5 - 9 μl) were put into triplicate wells of each respective primer in the same respective 96-well plate for all experiments. Primers for *Blos1* and GAPDH were used (Table S2). The cDNA and master mix were transferred to a 384-well plate using E1 ClipTip pipettor (4672040 Thermo Scientific). The qPR-PCR was run on a CFX384™ Real-Time System (Bio-Rad).

### RNA-seq analysis

All RNA-seq reads were mapped to *Mus musculus* reference genome GRCm38.p4, release 84, which is provided by Ensembl.org, by using STAR-2.5.2a (Dobin *et al*, 2012). The aligned reads were then counted by using RSEM-1.2.29 (Li & Dewey, 2011). Differential Expression Analysis was performed by using DESeq-1.30.0 (Anders & Huber, 2010) and edgeR-3.26.6 (McCarthy *et al*, 2012; Robinson *et al*, 2010). Differentially expressed gene (DEGs) was determined if the gene’s p-value (significance of differential expression) < 0.05 and the absolute value of the fold change > 1.5.

### Bioinformatic analysis

KEGG enrichment pathway analysis was performed by using WebGestalt-2013 (WEB-based Gene Set Analysis Toolkit) (Wang *et al*, 2013) with RIDD genes. Gene Ontology (GO) Enrichment was performed by feeding the significantly differentially expressed genes (DEGs) to PANTHER-v7 (Gaudet *et al*, 2011; Mi *et al*, 2010) in all three classes: Molecular Function, Biological Process and Cellular Component. Both KEGG pathway and GO enrichment analysis were filtered and sorted by Fisher test and Benjamini & Hochberg adjustment. The GeneCards website (Stelzer *et al*, 2016) was used to connect to the REFSEQ mRNAs from NCBI’s GenBank. The FASTA option was chosen and the fasta file was saved. The fasta file was uploaded to RNAfold Web Server (Institute for Theoretical Chemistry at University of Vienna) with the fold algorithm options of minimum free energy and partition function and avoid isolated base pairs. The Forna option was then used to view the secondary structure and manually searched for a stem loop with NNNNNCNGNNGNNNNNN. Interaction network analysis of KEGG pathways and the RIDD genes (p < 0.05) identified in this work was visualized using Cytoscape (https://cytoscape.org/) as described previously (Zhao *et al*, 2017).

### Generation of non-functional and RIDD resistant *Blos1* variants in RAW264.7 murine macrophages

Non-functional *Blos1* variants in RAW264.7 murine macrophages were generated using a protocol previously described (Hoffpauir *et al*, 2020). One clone containing either an amino acid deletion substitution or deletion in one of the XAT regions of murine BLOS1 (**Figure S4E**) was selected. For generation of Rr-*Blos1* variant and its control (wt-*Blos1*) in RAW264.7 murine macrophages, RNA was first extracted using RNeasy Plus mini-kit. *Blos1* cDNA was generated from mRNA and cloned into pCR™ 2.1-TOPO vector. Site-directed mutagenesis was utilized to generate a g449t mutation into one of the *Blos1* plasmid clones (**Figure S4F**). Both the wild-type (WT) and mutated *Blos1* segments were removed from the pCR™ 2.1-TOPO vector and cloned into pE2n vectors. Gateway cloning was used to generate expression vectors pLenti-CMV-Hygro-DEST (w117-1)-Blos1-WT and pLenti-CMV-Hygro-DEST (w117-1)-Blos1-G449T. At every step post cDNA creation, plasmid amplification vectors were verified via sequencing. The plasmids were then transfected into Lenti-X cells with psPAX2 and VSVG packaging plasmids. The virus was collected at 24 and 48 hrs post-transfection and stored at -80°C. RAW264.7 cells were transduced with the virus and cells containing the wt- or Rr-*Blos1* expression cassette were selected using hygromycin B (500 μg/ml).

### Drug treatments

Host cells were coincubated in 24-well plates with tunicamycin or 4μ8C at the indicated concentrations. Cells were treated with drugs 1 hr before, and during, infection with the indicated *Brucella* strains and incubated at 37°C with 5% CO_2_. At the indicated time points post infection, the treated cells were fixed with 4% formaldehyde and stained for immunofluorescence analysis or lysed to perform CFU assay or for RNA extraction assays as described above. To investigate whether the drugs inhibit *Brucella* growth, the drugs were individually added to *Brucella* TSB cultures at 37°C and incubated for 1 and 72 hr. CFU plating was used to assess bacterial growth in the presence of drugs, and thereby to evaluate the potential inhibitory effects. Host cells in which drug treatment or *Brucella* infection induced no significant differences in viability and membrane permeability as well as drugs that have no adversary effect on *Brucella* growth were used in the experiments reported in this work.

### Confocal immunofluorescence microscopy assays

Immunofluorescence microscopy staining and imaging methods to determine *Brucella* intracellular trafficking and colocalization of the bacteria and host BORC components in infected host cells were perform as previously described (Pandey *et al*, 2017; Pandey *et al*., 2018; Qin *et al*, 2011; Qin *et al*., 2008) with minor modifications. Briefly, to visualize Bm16M intracellular trafficking, the indicated host cells (5.0 × 10^4^ for 24-well plates and 1 × 10^4^ for 96-well plates) were seeded on 12-mm coverslips placed on the bottom of wells of 24-well plates or 96-well plates (without coverslip) and infected with Bm16M-GFP or Bm16M. At 0.5 (for 24-well plates) or 1 (for 96-well plates) h.p.i., the infected cells were washed with 1 × PBS and fresh media containing 50 μg/ml gentamicin was added to kill extracellular bacteria. At the indicated time points post infection, the infected cells were fixed with 4% formaldehyde at 4°C for overnight or for 20 minutes at 37°C before confocal fluorescence or immunofluorescence microscopy analysis was performed as previously described (Pandey *et al*., 2017; Pandey *et al*., 2018; Qin *et al*., 2011; Qin *et al*., 2008). The primary antibodies used were listed in the Key Resources Table. Samples were stained with Alexa Fluor 488-conjugated, Alexa Fluor 594-conjugated, and/or Alexa Fluor 647 secondary antibody (Invitrogen/Molecular Probes, 1:1,000). Acquisition of confocal images, and image processing and analyses were performed as previously described (Pandey *et al*., 2017; Pandey *et al*., 2018; Qin *et al*., 2011; Qin *et al*., 2008).

The BioTek Cytation 5 and Gen 5 software (version 3.05) were used to calculate perinuclear Lamp-1 index and autophagic flux. Specifically, for perinuclear Lamp-1 index, the area of the nucleus and average intensity for Lamp1 was measured, noted as A_N_ and Int_N_, respectively. The area of the whole cell and the average intensity for Lamp1 was measured, noted as A_WC_ and Int_WC_, respectively. Next the area of the cell 3 micrometers off from the nucleus and the average intensity for Lamp1 was measured, noted as A_C_ and Int_C_, respectively. Then, Int_WCN_

(Average intensity of Lamp1 in the whole cell minus nucleus) and Int_P_ (Average intensity of Lamp1 in the perinuclear region) were calculated as the following formula:

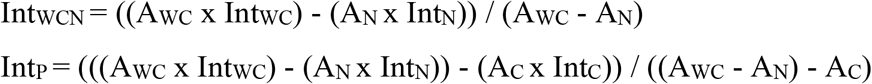

If Int_P_/Int_WCN_ ≤ 1, which means that there is no or very little perinuclear colocalization; If Int_P_/Int_WCN_ > 1, which means that there is perinuclear colocalization.

For autophagic activity, using BioTek Cytation 5 and Gen 5 software (version 3.05), the mean intensity of LC3b in the nucleus (M1) and in the cytoplasm (M2) were measured using a primary and secondary mask in every individual cell. In the Gen 5 software, a subpopulation analysis was carried out to identify cells that had a ratio of M2/M1 <1. From this, the perinuclear LC3b index or autophagic activity was calculated as the following formula: Perinuclear LC3b index (or autophagic activity) = (Total number of identified cells with M2/M1 <1)/Total number of the analyzed cells.

For calculation of the BCV-BLOS1^+^ index, the corrected total cell fluorescence was measured by taking the integrated density (area of cells × mean fluorescence), then the number of *Brucella* in each cell and colocalization of BLOS1 and Bm16M (BCV-BLOS1^+^, %) were counted. Cells that did not contain bacteria were removed from the calculation. The integrated density was divided by the number of bacteria in the cell to obtain the “Total fluorescence per bacteria”. The “BCV-BLOS1^+^ index” was then calculated by multiplying the “Total fluorescence per bacteria” with the colocalization percentage. Since BLOS1 protein is more stable in control cells treated with 4μ8C, the value of BCV-BLOS1^+^ index was normalized as 100%.

### Protein pull-down assays and immunoblotting analysis

Pull-down assays for testing physical interaction of proteins were perform using the Pierce Co-Immunoprecipitation (Co-IP) Kit (Thermo Scientific, USA) according to the manufacturer’s instructions. Preparation of protein samples and immunoblotting blot analysis were performed as described previously (Ding *et al*, 2021; Pandey *et al*., 2017; Pandey *et al*., 2018; Qin *et al*., 2011). Densitometry of blots was performed using the ImageJ (http://rsbweb.nih.gov/ij/) software package. All Westerns were performed in triplicate and representative findings are shown.

### Statistical analysis

All the quantitative data represent the mean ± standard error of mean (SEM) from at least three biologically independent experiments, unless otherwise indicated. The data from controls were normalized as 1 or 100% to easily compare results from different independent experiments. The significance of the data was assessed using the Student’s *t* test (for two experimental groups), 2-way ANOVA test with Holm-Sidak’s multiple comparisons, or the Kruskal-Wallis test with Dunn’s multiple comparison. For the RNA-seq results, Log_2_ fold changes were calculated and results were screened to meet the threshold (| log_2_FC (fold change) | > 1, P < 0.05) for selection. DEGs met the criteria and in UPRsome were included in the final lists.

## Key resources

All key resources, including bacterial strains, mammalian cell lines, reagents, etc. used in this work are listed in Table S3.

## Data availability

All data are included in this article, Supplemental Figures and Supplemental Datasets. Source Data files have been provided for Figures 1,4 and 7.

**Expanded view** for this article is available online.

## Author Contributions

P.dF., K.M.W. and Q.M.Q. conceived and designed the experiments; K.M.W., Q.M.Q., A.P., A.C., D.Z., J.Y., G.G., F.L., A.L.C., H.Q.F. performed the experiments; K.H., S-H.S., X.Q., P.dF., Q.M.Q., K.M.W. Y.L., H.W.C., X.Q.L., H.Z., A.P., and J.L. analyzed the data: P.dF., T.A.F., Q.M.Q., A.R.F., K.H., S-H.S., X.Q., H.W.C., and H.Z. R.P.M., C.D.J., L.B., K.P. and J.L. contributed reagents/materials/analysis tools; P.dF., T.A.F. and Q.M.Q. supervised the work; Q.M.Q., P.dF. and K.M.W. wrote the manuscript.

## Acknowledgements

The authors are grateful to Dr. Julie Hollien (University of Utah) for sharing key cell lines used in this study, to Drs. Christine McFarland, Jessica Bourquin, Todd Wisner and Susan Gater (Texas A&M) for key support, and to Drs Steve Fullwood and Kalika Landua (Nikon Instruments) for expert assistance with the microscopy analysis. This work is supported by the Texas A&M Clinical Science Translational Research Institute Pilot Grant CSTR2016-1, DARPA (HR001118A0025-FoF-FP-006), NIH (R21AI139738-01A1, 1R01AI141607-01A1, 1R21GM132705-01), the National Science Foundation (DBI 1532188, NSF0854684), the Bill Melinda Gates Foundation, and the Defense Advanced Research Projects Agency (Agreement HR001118A0025-FoF-FP-006) to PdF; the National Natural Science Foundation of China (# 81371773) to QMQ. The National Institute of Child Health and Human Development [RHD084339] to TAF (in part). The National Science Foundation Grant 1553281 to XQ. The content of the information does not necessarily reflect the position or the policy of the Government, and no official endorsement should be inferred.

